# KATK: fast genotyping of rare variants directly from unmapped sequencing reads

**DOI:** 10.1101/2020.12.23.424124

**Authors:** Lauris Kaplinski, Märt Möls, Tarmo Puurand, Fanny-Dhelia Pajuste, Maido Remm

## Abstract

**Motivation:** KATK is a fast and accurate software tool for calling variants directly from raw NGS reads. It uses predefined k-mers to retrieve only the reads of interest from the FASTQ file and calls genotypes by aligning retrieved reads locally. KATK does not use data about known polymorphisms and has NC (No Call) as default genotype. The reference or variant allele is called only if there is sufficient evidence for their presence in data. Thus it is not biased against rare variants or *de novo* mutations.

**Results:** With simulated datasets, we achieved a false negative rate of 0.23% (sensitivity 99.77%) and a false discovery rate of 0.19%. Calling all human exonic regions with KATK requires 1-2 h, depending on sequencing coverage.

**Availability:** KATK is distributed under the terms of GNU GPL v3. The k-mer databases are distributed under the Creative Commons CC BY-NC-SA license. The source code is available at GitHub as part of Genometester4 package (https://github.com/bioinfo-ut/GenomeTester4/). The binaries of KATK package and k-mer databases described in the current paper are available on http://bioinfo.ut.ee/KATK/.

## Introduction

### Need

A typical personal genome differs from the reference human genome at 4 to 5 million positions (1000 Genomes Project Consortium *et al.*, 2015). Whole-genome sequencing (WGS) and whole-exome sequencing (WES) allow systematic discovery and description of these differences in personal genomes (Gudbjartsson *et al.*, 2015; Telenti *et al.*, 2016; Lek *et al.*, 2016; Cirulli *et al.*, 2020; Locke *et al.*, 2019; Lohmueller *et al.*, 2013). Many of these variants are likely to have functional implications. For example, human genomes contain approximately 100 loss-of-function (LoF) variants with ~20 genes completely inactivated (MacArthur *et al.*, 2012). Variant detection with WGS and WES is widely used in medical genomics (Daneshjou *et al.*, 2017), particularly in the diagnostics of rare diseases (Danielsson *et al.*, 2014; Sawyer *et al.*, 2016; Wright *et al.*, 2018) and cancer genomics (Berger and Mardis, 2018; Nakagawa and Fujita, 2018; Xu *et al.*, 2019). In this article, we focus on the analysis of WGS data only, as it provides more uniform coverage of all genomic regions than WES.

### Speed

Variants in the human genome are typically genotyped by mapping sequenced reads and then performing genotype calling (Langmead and Salzberg, 2012; McKenna *et al.*, 2010; Li and Durbin, 2010; Li and Homer, 2010). A standard pipeline requires 40-50 h to process a high-coverage human genome from raw WGS data to variant calls on a multithreaded server. Medical uses of WGS are often time-critical (Mestek-Boukhibar *et al.*, 2018; Buchan *et al.*, 2019). Typically, attempts to reduce the time spent processing a single genome focus on the parallelization or optimization of existing algorithms (Rimmer *et al.*, 2014; Freed *et al.*, 2017; Kim *et al.*, 2018; Poplin *et al.*, 2018).

### Alignment-free variant calling

Another approach for accelerating the processing of genome data is to use alignment-free algorithms for calling genotypes. This approach would eliminate the need for mapping and sorting the reads. Kimura and Koike introduced an alignment-free method that uses depth of coverage of variable length k-mers for detecting variants in human genomes. This method can handle WGS data in 15 h and WES data in 2.5 h (Kimura and Koike, 2015). Other examples of alignment-free variant calling software are COBASI (Gómez-Romero *et al.*, 2018), Kestrel (Audano *et al.*, 2018), LAVA (Shajii *et al.*, 2016) and FastGT (Pajuste *et al.*, 2017). COBASI utilizes 30-mers to analyse unique regions of the genome (84% of the genome) from WGS data in 31.5 h. WES data could be analysed in less than 9 h, although this was not directly tested. These researchers achieved 96% AUPR (Area Under Precision-Recall Curve) with simulated data (Gómez-Romero *et al.*, 2018). Kestrel is an alignment-free variant caller that uses haplotype reconstruction from k-mer frequencies and is capable of characterizing densely packed SNPs and large indels. However, it has been tested on bacterial genomes only, not on human WGS or WES data(Audano *et al.*, 2018). LAVA is able to predict the genotypes of known SNVs from WGS data in 5 h with 93% accuracy (Shajii *et al.*, 2016). FastGT is able to call SNV genotypes of known variants from WGS data in 20 minutes with 99.6% accuracy (Pajuste *et al.*, 2017). However, both FastGT and LAVA are limited to the analysis of previously known variants.

### Reliability of calls

Previous studies indicate that a significant fraction of the human genome is accessible to analysis with short k-mers. For example, 85.7% of the human genome is covered with uniquely located exact 32-mer matches, and 79.3% of the human genome is covered with uniquely located 32-mer matches with no Hamming neighbours (Shajii *et al.*, 2016). Telenti and others have demonstrated that 84% of an individual human genome can be called confidently from deep sequencing data. This high-confidence region includes 91.5% of exons (Telenti *et al.*, 2016). We have shown previously that alignment-free methods can be used to reliably genotype 74% of known SNVs (Pajuste *et al.*, 2017) and 77% of polymorphic Alu element insertions (Puurand *et al.*, 2019). It is important to note that inconsistent genotype calls are associated with certain inaccessible genomic regions (Derrien *et al.*, 2012; Lee and Schatz, 2012; Weisenfeld *et al.*, 2014), whereas variant calling in the remaining 75-85% of the genome is robust and reliable, even with short k-mers.

Traditional variant calling pipelines that use the information from full-length paired-end reads typically attempt to call the entire genome, assuming that most reads are uniquely mappable. In reality, the “inaccessible” genomic regions remain error-prone even for paired-end reads. For example, it has been observed that different variant calling pipelines generate slightly different genotype calls from the same data (O’Rawe *et al.*, 2013; Pirooznia *et al.*, 2014; Li, 2014; Sandmann *et al.*, 2017). This results in a mix of reliable and unreliable variant calls from “inaccessible” regions. Often, reliable and unreliable calls are not clearly distinguished in software output.

Ultimately, the variant calling software should clearly distinguish reliable calls from unreliable calls and at the same time should be as fast as possible. With these requirements in mind, we developed a new and original alignment-free method, KATK, which is focused on the fast and reliable analysis of accessible regions of the genome. KATK identifies unreliable variant calls and clearly distinguishes them in the output. To demonstrate the capabilities of the KATK algorithm, we compiled a set of k-mers from human exonic regions and tested the performance accuracy of the algorithm on various datasets.

## Methods

KATK detects sequence variants by aligning selected reads to short segments (140 bp) of the reference genome. The relevant reads are obtained from full sequencing data by unique tag k-mers. By requiring uniqueness, KATK avoids time-consuming and error-prone mapping of reads to regions of the genome that are not sufficiently unique. Although many regions of the genome remain uncallable by this method, the remaining ones can be called faster and with high accuracy. The algorithm is deterministic and trivially parallelizable.

KATK uses a hypothesis-free approach for calling each mutation site, i.e. the knowledge about known polymorphisms in population and their allele frequencies is not used.

The KATK algorithm relies on a set of k-mers that are unique in a given genome. There are no specific requirements for the k-mers, except that they must have the same fixed length and must be uniquely located in the reference genome.

The input for analysis of individual genomes is a FASTQ file with raw sequencing reads and a preprocessed database of k-mers (Figure 1). The k-mer database consists of a searchable tree of k-mer sequences in binary format (described in (Pajuste *et al.*, 2017)) and a text file with the description of genomic regions of interest. The more detailed description of database files is given in Supplementary File 1. To minimize the noise from sequencing errors, tag k-mers are selected to have as big Hamming distance with the next closest k-mer from any other region of the genome as possible. To reduce the bias caused by partial alignment at region ends and streamline processing, the long regions are broken into overlapping 140-bp-long chunks with a 50-bp step. The database used in the current work is downloadable from http://bioinfo.ut.ee/KATK/downloads/.

**Figure 1.**
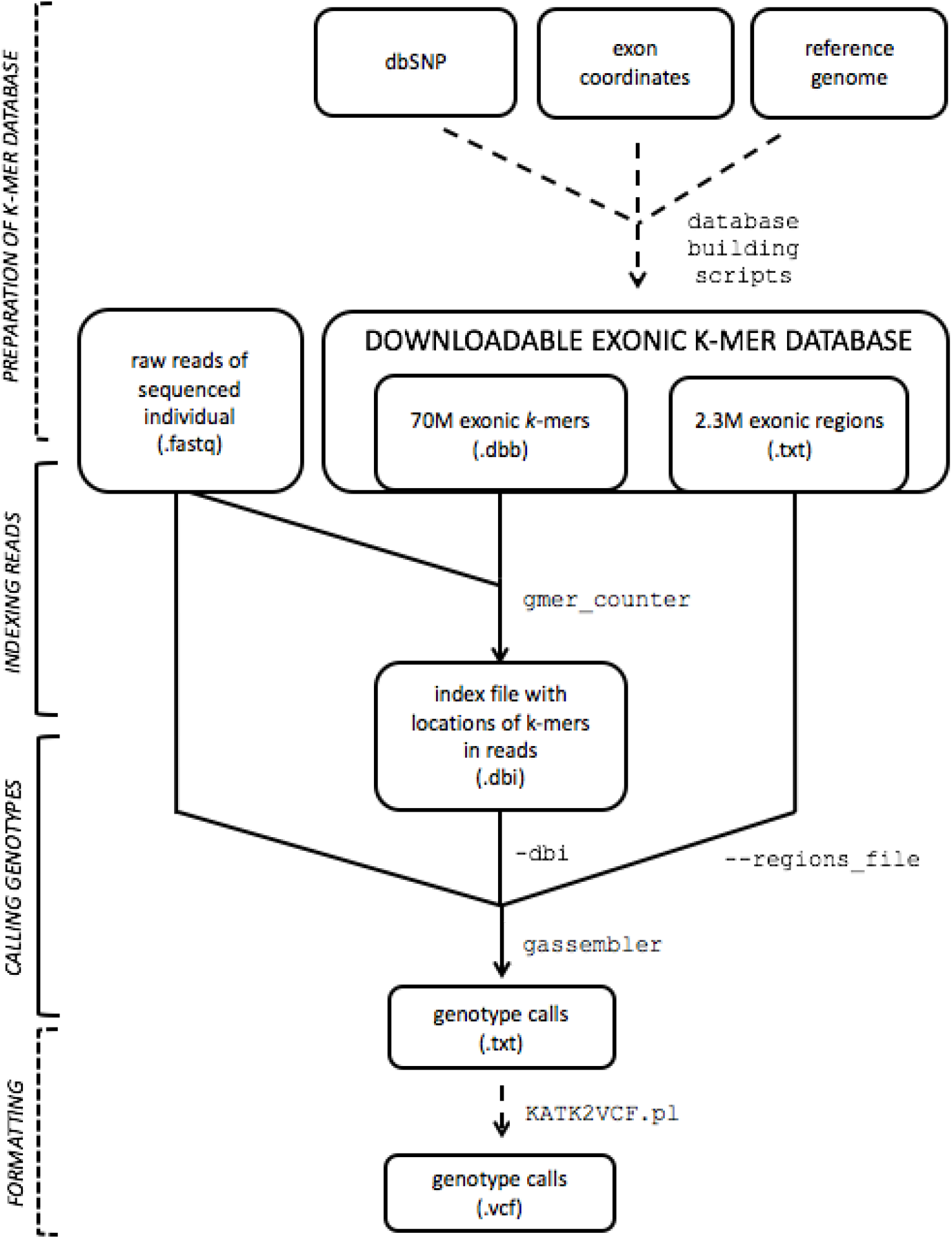
Description of different steps and data for alignment-free calling of rare variants.

For analysis of each individual genome, two separate steps are executed. In the first step, the sequencing reads containing each k-mer are written to an index file. In the second step the chunks are processed one-by-one. The reads containing at least one specific k-mer of a given chunk are retrieved from the FASTQ file using the index. Reads are locally aligned to the reference sequence of the chunk and grouped by the haplotype. Genotypes are called based on allele counts of the best groups within each position of the alignment.

### Algorithm

For each analyzed segment, there is a corresponding list of unique tag 25-mers from 50 bp before to the start to 50 bp after the end of the segment in regions file.

For each segment, a list of reads containing at least one of these tag 25-mers is obtained. Reads are then aligned to the reference segment with the Smith-Waterman algorithm, and those with too short or divergent alignment are discarded.

Then, a list of all positions where at least 2 reads have an identical difference from the reference is composed. These positions are potential mutation sites in a given individual.

Next, reads are grouped by a greedy algorithm such that all reads in the same group have either identical or compatible alleles for each potential mutation site. Groups are ordered by similarity to the reference genome. For haploid chromosomes, the one best group is selected, for diploid chromosomes either one or two groups. The consensus alignment of each group corresponds to a haplotype of a given region in a sequenced individual.

As the segments overlap, for any single position, there may be multiple calls. The best call is selected by the selection model, trained on simulated individuals. The call quality is then estimated based on one of the two quality models (logistic regression models; estimated using data from simulated individuals).

Model 1 - the quality of non-reference calls (45 parameters)
Model 2 - the quality of reference calls (21 parameters)

The meaning of model 1 is straightforward - a probability of making the wrong non-reference call with given alignment parameters. The positions with a quality score below the cutoff are marked as NC.

The meaning of model 2 is different. As this model is applied to reference calls, it estimates whether the algorithm would have detected the mutation if there had been one. Positions with a quality score above the cutoff are omitted from the output (considered to be the same as reference), and positions with a quality below the cutoff are marked as NC.

Model 2 uses as an input only information that would likely not be influenced by the true status (reference/non-reference) of the genome. Hence, model 2 uses fewer parameters than model 1. The full description of models is given in Supplementary File 2.

### Diploid regions

For diploid regions, if there is more than one group of reads, KATK has to make a decision whether to use one or two of these groups for calling. The selection is done separately for each potential mutation site by evaluating the probability of getting a given distribution of nucleotide counts from a multinomial distribution. The full formula used is given in Supplementary File 3.

### Haploid regions

For haploid regions (sex chromosomes in males and mitochondria), haploid versions of the models and different expected coverage are used. For sex chromosomes, the expected coverage is half of the autosome coverage; for the mitochondrial genome, the average number of mitochondrial reads in a given chunk is used.

## Implementation

### Program code

KATK is written in C as part of the GenomeTester4 package. It does not require external dependencies aside from libpthread and zlib that are standard parts of all Linux distributions.

### Preparation of the k-mer database

Exons were defined as protein-coding regions of the genes plus 50 bp flanking region from both sides of the region. Corresponding genomic coordinates used in this study were extracted from the Ensembl Biomart database (Ensembl Release 89). The sequences of the exons were created based on these coordinates and the reference genome (GRCh38p10). Overlapping exon sequences were combined. Flanking regions of 12 bp were added to both ends of the combined regions allowing detection of the first nucleotide of the region by 25-mers.

A set of validated biallelic SNVs was also extracted from Ensembl Biomart. For every 25-mer that contained less than three SNVs, possible alternative sequences were created; thus, depending on the number of SNVs (0, 1 or 2), the database contained either 1, 2 or 4 k-mers for every location in the exome.

### Compromise between call rate and accuracy

Two probabilistic models are used for calling genotypes (described in Methods). One model tests the likelihood of the presence of alternative allele P_mut_, and the other estimates the probability of discovering a difference from the reference in a particular position (if there actually would be a difference) P_ref_. If both models give low likelihood, we assign the position as the NC (No Call) position. In other words, the default call is No Call, unless we have sufficient data to support the genotype.

In a typical classification test, one has to make compromises between specificity and sensitivity. Increased sensitivity normally results in decreased specificity and vice versa. In our setup, the compromise is between call rate and accuracy. This reduces the number of false negative calls, particularly in cases where the depth of coverage is low.

Our approach influences the procedure for the evaluation of calls. In most cases, the variant calling methods are evaluated based on what fraction of variants they are able to detect from all variants in the genome. In our case, we use only callable positions in evaluations (Figure 2).

**Figure 2.**
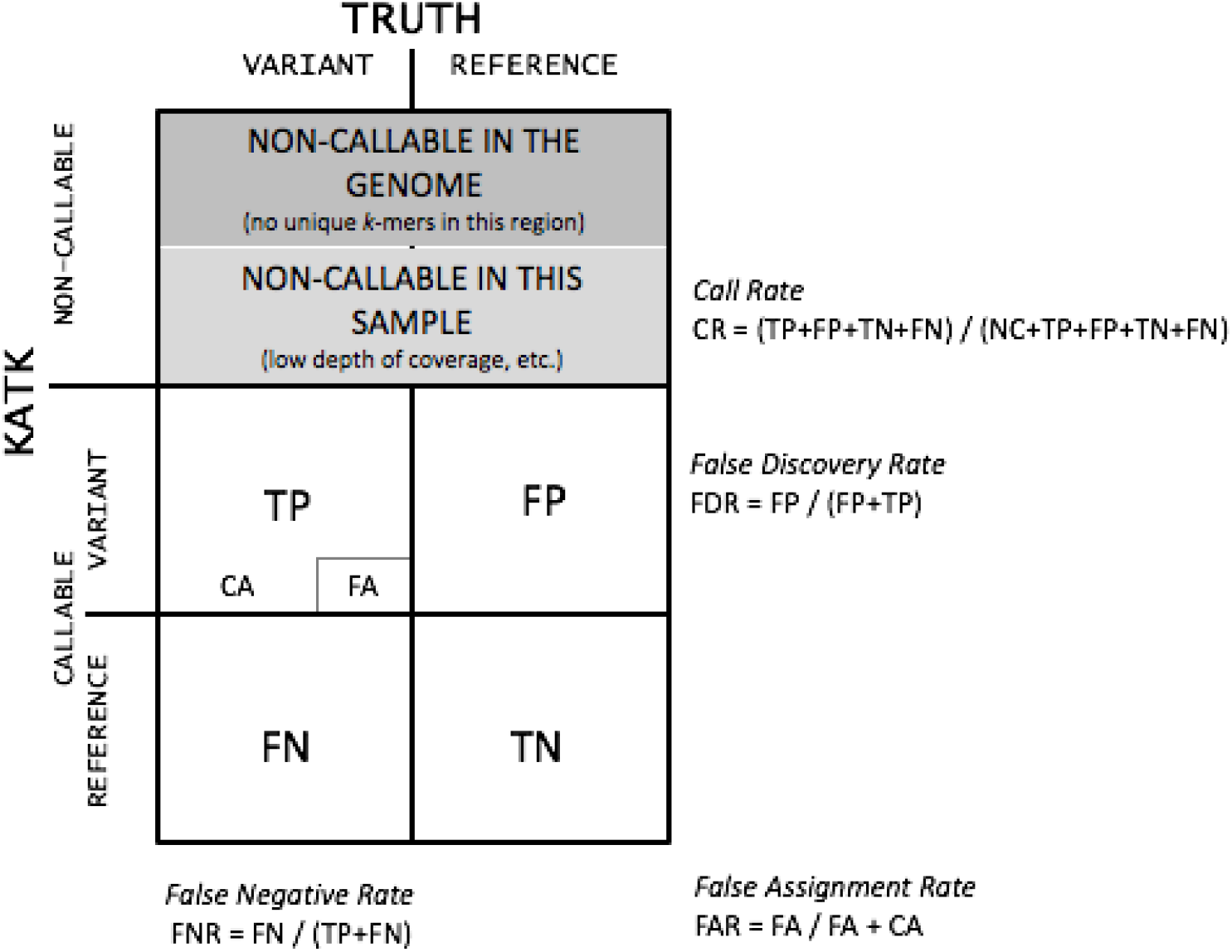
Explanation of terminology used in the evaluation of KATK. The main characteristics are the call rate (CR), false discovery rate (FDR) and false negative rate (FNR). FA denotes false genotype assignments, i.e., cases where a variant was detected but was assigned an incorrect genotype. CA denotes the correct genotype assignment. FA and CA are subsets of TP.

The accuracy can be adjusted by the quality cutoffs P_mut_ and P_ref_. A higher value of P_mut_ provides higher accuracy (both FDR and FNR are lower) but a lower call rate (Figure 3 C and D). Reducing the value of P_mut_ increases the overall call rate but reduces the accuracy of calls. A higher value of P_ref_ makes KATK more conservative for calling reference alleles (reduces both FNR and call rate). This property enables users to adjust the stringency of analysis. The user can adjust P_mut_ and P_ref_ with a command-line parameter, and we used P_mut_=0.95 and P_ref_=0.5 throughout this article.

**Figure 3.**
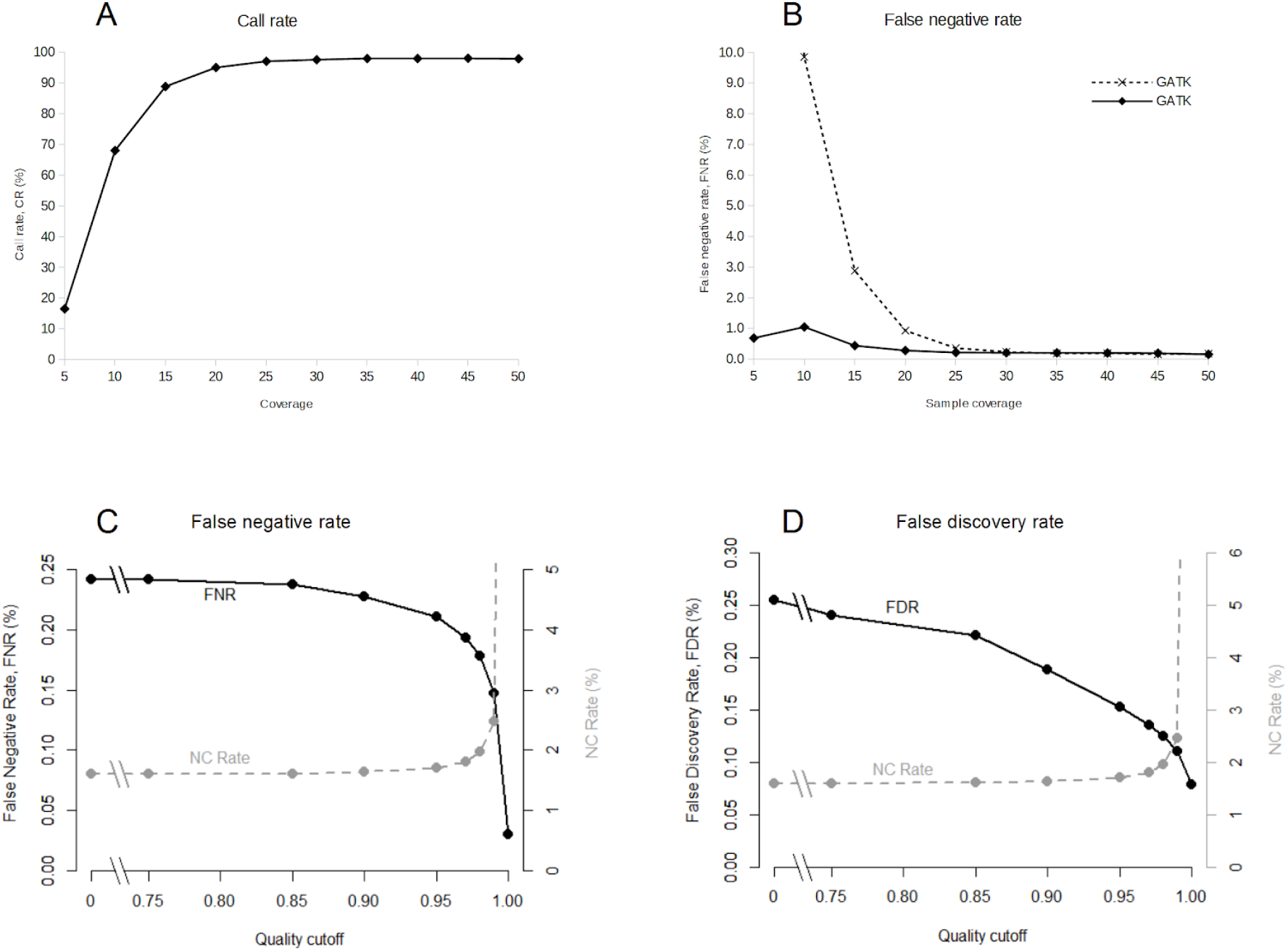
Effect of sample coverage and quality cutoff on KATK sensitivity and selectivity. A) The effect of depth of coverage on the call rate (CR) on simulated data. The probability cutoff of calls was set to 0.95 and of reference positions 0.5. B) The effect of the depth of coverage on the false negative rate for KATK and GATK. Simulated datasets were used. For KATK, the probability cutoff of calls was 0.95 and that of reference positions was 0.5. C) The effect of ‘quality cutoff’ on false negative rate (FNR) and no call rate (NCR). D) The effect of ‘quality cutoff’ on false discovery rate (FDR) and no call rate (NCR).

## Validation

We tested the program with simulated data and two high-quality real datasets. The simulation method is described in Supplementary File 4.

### Effect of depth of coverage on call rate and accuracy

The effect of sequencing coverage and the performance of the program was tested on simulated reads. Coverage ranged from 5x to 50x with step 5. Two simulations were created for each coverage - one female and one male.

KATK requires at least two confirming reads, one from either direction, to call a mutation. It is thus expected that at low sequencing depth, many mutations remain undetected because of stochastic variability in coverage (Figure 3 A). Thus, we discourage using KATK calling for sequence depths less than 20. A depth above 30 does not significantly improve the call rate. At 30x coverage, KATK had a sensitivity of 99.77% and a false discovery rate of 0.19%.

By using two separate models, KATK is able to distinguish the situations where true reference alleles are detected and where the decision cannot be made. The majority of low-coverage regions will be marked as uncalled (NC) instead of reference alleles. This results in the capping of false negatives at approximately 1%, even for low sequencing depth (Figure 3 B). Thus, the main effect of low sequencing depth for KATK is the falling call rate and not the increase in false negatives.

### Reference datasets

We tested the accuracy of KATK on Illumina Platinum individual NA12878(Eberle *et al.*, 2017), sequenced to 50x coverage. The published sequence of a given individual is derived from variant calls with different methods, adjusted by the reference data set of 5.4 million phased human variants validated by genetic inheritance from sequencing a three-generation 17-member pedigree.

Additional accuracy testing was performed by analysing published sequencing reads from fully homozygous cell lines CHM1 (complete hydatidiform mole) and CHM13 and their 1:1 mixture (synthetic diploid genome)(Li *et al.*, 2018).

### Concordance of SNV and indel calls on the Platinum dataset

KATK has high concordance (98.4%) with SNV genotypes in the Illumina Platinum (Eberle *et al.*, 2017) dataset (Table 1). Indel calling with KATK is clearly less accurate than SNV calling. Single nucleotide insertions and deletions are 91% concordant with Illumina Platinum calls. Longer insertions are detected at a slightly better rate with 94.1% concordance. However, long deletions have a high frequency of false negatives (11.6%) and false positives (22.7%).

**Table 1.** Concordance of variants (in percent) with the Platinum dataset, generated from CEPH individual NA12878. SNV - Single nucleotide variants; INS1 - insertions of a single nucleotide; INS>1 - insertions of more than one nucleotide; DEL1 - deletions of a single nucleotide; DEL>1 - deletions of more than one nucleotide.

|  | SNV | INS1 | INS>1 | DEL1 | DEL>1 |
| --- | --- | --- | --- | --- | --- |
| FDR | 2,8 | 4,2 | 3,7 | 4,9 | 22,7 |
| FNR | 1,6 | 7,2 | 5,9 | 9,0 | 11,6 |

### Mendelian errors on the CHM dataset

The CHM dataset (Li *et al.*, 2018) provides us with an independent method for verifying calling accuracy in the real sequencing data because all true mutations have to adhere to a simple rule: be homozygous in “parents” and have a Mendelian pattern in “child”. Although systematic errors cannot be detected in this way, non-systematic errors are detected with high probability. We analysed all three sequencing samples with both GATK (McKenna *et al.*, 2010) and KATK and counted concordance and divergence from the expected pattern for each called mutation. The results are shown in Table 2. Both methods have positions where detected mutation patterns are not in concordance with known genotype models.

**Table 2.** Frequency of different non-mendelian error scenarios in CHM trio dataset calls by GATK and KATK. “+” denotes cases where both “parents” were homozygous and “child” has an expected genotype. “−” denotes cases where either at least one “parent” was heterozygous or “child” unexpected genotype, not concordant with Mendelian inheritance.

| GATK | KATK | Frequency |
| --- | --- | --- |
| + | + | 83,4% |
| + | - | 8,9% |
| - | + | 5,6% |
| - | - | 2,1% |

### Computational performance

We tested KATK performance on a CentOS workstation with 32 2.0GHz CPUs and 512 GB of RAM. The maximum number of concurrent threads was set to 24. For normal sequencing depth, the full calling pipeline took approximately 1 h (Figure 4).

**Figure 4.**
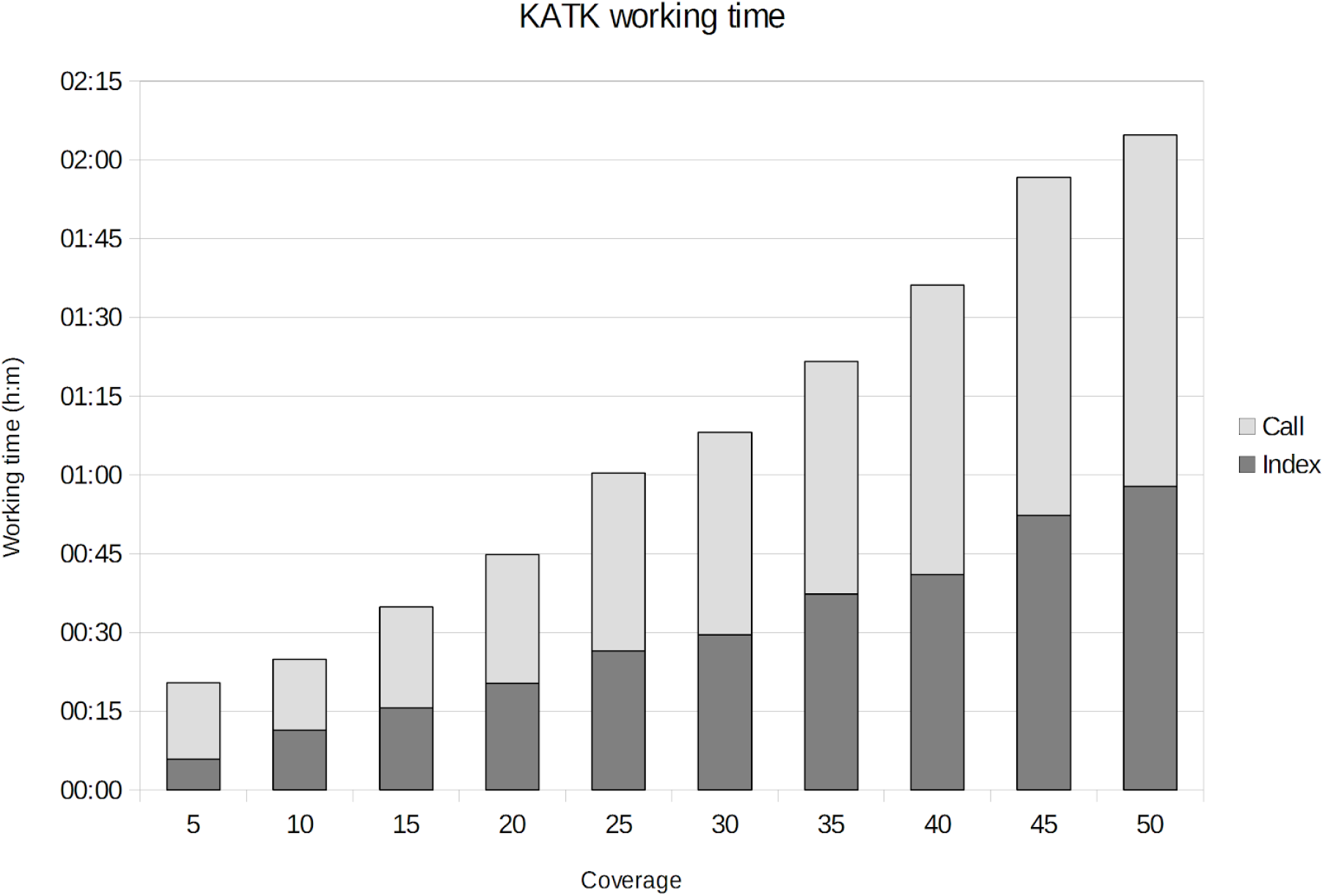
The computation time of KATK. The computation time of both indexing and calling grows linearly with the size of the FASTQ file. The tests were conducted on a CentOS 5.10 Linux server with 32 cores (2.27 GHz) and 512 GiB (gibibyte, 2^30^ bytes) RAM, and the maximum number of threads was set to 24.

KATK uses memory mapping of files with prefetching to increase performance. Thus, the maximum resident memory consumption is determined by the size of FASTQ files. It can run on a workstation with significantly less memory but with reduced performance. For optimal performance, at least 200 GB of RAM is needed for 25x coverage and 300 GB for 50x coverage.

## Discussion

Although the popularity of long-read WGS technologies is on the rise, short-read technologies remain viable as a cost-effective alternative for genome analysis. Short read sequencing combined with alignment-free (mapping-free) analysis methods can offer significantly faster throughput of large datasets. KATK is able to call exonic variants from WGS data in less than 2 h. Furthermore, KATK is able to estimate the reliability of calls and output only those variants that exceed a certain level of confidence. As a completely original algorithm, KATK complements traditional calling algorithms, detecting variants that otherwise might remain unnoticed (Table 2).

### Hypothesis-free approach

Although taking existing knowledge about polymorphisms and their allele frequencies into account or doing joint calling of many individuals increases the precision of calls, in some cases the joint calling might not be practicals. We designed KATK to be hypothesis-free - i.e. it considers each nucleotide in reference sequence having an equal probability of mutation. Thus KATK is best suited for calling previously unknown (rare or de-novo) variants. KATK can be complemented with SNV genotyping tools, e.g. FastGT, to increase the reliability of the calls of previously known polymorphisms. In future, we plan to develop a new version of KATK that allows joint calling of multiple individuals.

### Length of the k-mer

KATK is optimized to work with short k-mers. In this article, we use 25-mers, but any k-mer with a length between 22 and 32 would give similar results (Pajuste *et al.*, 2017). Longer k-mers have lower effective depth of coverage due to the edge effects in relatively short Illumina reads. Shorter k-mers might be less unique; therefore, the fraction of accessible regions of the genome would be lower.

### Analysing WES data

While WGS has the advantage of capturing all possible mutations, whole-exome sequencing (WES) still remains a cost-effective alternative that captures genomic regions with most phenotype-altering mutations. Unfortunately, enrichment or capture required for WES introduces high variation in depth of coverage that seriously hinders subsequent detection of variants. It has been shown that only 72 - 80% of calls from WES data are concordant between different technologies (Samorodnitsky *et al.*, 2015). The KATK genotype calling algorithm is particularly vulnerable to unpredictable fluctuations of coverage because the probabilistic models we use are based on even coverage for the entire genome. We tested KATK with simulated WES data and achieved FNR (18%) and FDR (10%). Although these rates are similar to other tools using WES data, we do not recommend the use of WES data with KATK.

### Possibilities for analysing a larger number of variants

In the current analysis, we used high coverage WGS data but called only exome regions, overall accounting for approximately 3% of the genome. The KATK algorithm is able to process arbitrary regions, the only requirement being the existence of unique tag k-mers. However, as the computation time is roughly proportional to the length of the analysed sequence, the speed advantage of KATK might be lost.

### Accuracy considerations

In the clinical context, both the presence and absence of mutations can be equally important for diagnosis. Therefore, both high sensitivity and high positive predictive value (PPV) are required. KATK performs well with simulated data (0.25% FNR) and with the Illumina Platinum dataset (1.6% FNR) for SNV calls. This result is comparable to mapping-based rare variant detection methods. Most methods demonstrate FNR between 1% and 2% in an unbiased CHM validation dataset (Li *et al.*, 2018).

Indel calling has a significantly lower accuracy than SNV calls. This result is comparable to the indel calling accuracy reported by other authors (Telenti et al. 2016; Li et al. 2018; Hasan et al. 2015).

It should be kept in mind that all methods for calling rare variants, either mapping-based or alignment-free, make significant amounts of errors and are not usable in medical diagnostics without additional validation steps. Sanger sequencing of variants of interest is still needed for KATK, as well as for other variant calling methods (Mu *et al.*, 2016).

## Supporting information

Supplementary text 1: Database overview

Supplementary file 3: Model descriptions

Supplementary file 4: Diploid model

Supplementary text 2: Simulation parameters

## Acknowledgements

This work was funded by institutional grant IUT34-11 from the Estonian Ministry of Education and Research and the EU ERDF grant No. 2014-2020.4.01.15-0012 (Estonian Centre of Excellence in Genomics and Translational Medicine). The cost of the WGS sequencing of the individuals from the Estonian Genome Centre was partly covered by the Broad Institute (MA, USA) and the PerMed I project. The genome data was collected and used with ethical approval Nr. 206T4 obtained for the project SP1GVARENG. The authors thank Mihkel Vaher for the constructive reading of the manuscript.

## Supplementary materials

**Supplementary file 1: Database**

**Supplementary file 2: Simulated genomes**

**Supplementary file 3: Model descriptions**

**Supplementary file 4: Discrimination between homozygous and heterozygous diploid calls**

