## Supplementary text 1: Database overview for "KATK: fast genotyping of rare variants directly from unmapped sequencing reads"

### Supplementary file 1

### Database

Exons were defined as protein coding regions of the genes, the coordinates used in this study were extracted from Ensembl (Release 89) Biomart database Ensembl Genes 89 and dataset Human genes (GRCh38p10). The chosen parameters were Exon stable ID, Chromosome/scaffold name, Exon region start (bp), Exon region end (bp). Chosen filters were Gene type: protein coding and Chromsome/scaffold: 1-22, X, Y and MT. The total number or regions in the datatable was 569 768.

The Exon region start coordinate was decreased by 62 bp and the Exon region end coordinate was increased by 62 bp to create exomic regions with small flanking areas. All overlapping regions were combined, leaving a total number of 213 417 exomic regions. The exome sequences were created based on these coordinates and the reference genome (GRCh38p10). All the sequences were then divided into overlapping 25-mers.

A set of SNVs and indels was also extracted from Ensemble (Release 89) Biomart database Ensembl Variation 89 and dataset Human Short Variants (SNPs and indels excluding flagged variants) (GRCh38.p10). The chosen parameters were Variant Name, Chromosome/scaffold name, Chromosome/scaffold position start (bp), Chromosome/scaffold position end (bp) and Variant alleles. Chosen filters were Chromsome/scaffold: 1-22, X, Y and MT, Variant source: dbSNP and Variant supporting evidence: Only. The total number of rows was 95 493 065.

The results were filtered to remove some inconsistensies or unwanted variations (reference allele variants not matching the reference sequence, non-unique or incorrect locations, start/end positions not matching the length of variant, sequences containing N's etc). Indels were also filtered out and from the remaining SNVs, a set was chosen so that there would be no more that three SNVs in a 25 bp.region. The final set contained 63 637 010 SNVs.

For every 25-mer in the exome the possible alternative sequences were created, thus depending on the number of SNVs (0, 1 or 2), the database contained either 1, 2 or 4 k-mers for every location in the exome. The total number of occurrences in the exome was also marked for each k-mer as well as a maximum of two other locations where the k-mer can be found.

The layout of binary k-mer database and the creation of extended human reference genome is described in:

Pajuste,F.-D. et al. (2017) FastGT: an alignment-free method for calling common SNVs directly from raw sequencing reads. Sci. Rep., 7, 2537.

### Regions file

To create a regions file all exons in the genome (with flanking 62bp areas) was scanned with 50bp step, starting at position 50 after the start of an exon. For each step, all k-mers in the 125bp region before the current position and 125bp after the current position were obtained. Non-unique k-mers were discarded and from the remaining, the ones with the least number of matches with hamming distance 2 were selected. If there was at least one k-mer both in the left and right 125bp region, the current position was included in regions list.

### Alignment

A reference sequence from 70bp before until 70bp after the current position was saved in the regions file. The full 140bp region was then aligned independently with all reads obtained via tag k-mers but only the middle 120bp part of alignment was used for alignment quality estimation and grouping. Then the final 100bp middle region was used for calling. This reduced the effect of unreliable alignment at region endpoints, especially for insertions and deletions.

The region file used in current publication can be downloaded from:
http://bioinfo.ut.ee/KATK/downloads/
