## Supplementary file 3: Model descriptions for "KATK: fast genotyping of rare variants directly from unmapped sequencing reads"

**Supplementary file 2: Models**

Explanation

**Model 1a (call selection)**

Observations:

For all position (separately for each chunk) one or possibly two (heterozygous and homozygous) raw calls were generated based on the local alignment of the reads in the chunk.

These raw calls from all chunks were used as inputs to the logistic regression model (possibly containing many calls for the same position in genome). Binary variable indicating the correctness of the call was used as response variable (as we used simulated data to fit the model the true genotype was known to us). Variables like distance from the region boundary, coverage, call type etc were used as explanatory variables.

The model is later used for selecting the best call (over all chunks) for any particular position. The raw call achieving highest probability of being correct is selected as the best call for a given position.

**Model 1b (quality score for the best call)**

After selecting one call for each position (by using the Model 1a) another logistic regression model is fitted to the selected best calls dataset. Response variable is the correctness of the (selected best) call.

**Model 2 – the quality of reference calls**

The aim of the model: if there would have been an mutation in a given position, could we detect it? To estimate this model only non-reference positions were used. The y variable is a binary variable indicating wether the call is correct (True positive) or not correct (False negative or a wrong non-refernce call). Independent variables used contain only information about the position (distance from the region boundary) or about overall coverage.

Models were estimated on 20 sets of simulated sequencing reads (using the same simulation parameters but different mutations than the ones used for quality assessment). The coverages were from 5 to 50 with step of 5, one male and one female for each coverage.

The meaning of parameters is:

```
GLOBAL_COV: the average sequencing depth of the individual
COV: the number of reads covering given position (in final call, i.e. reads in
discarded groups are removed)
GRP: the number of groups for calling
COMP_2: the number of reads compatible with the first two groups.
G0_COMP: the number of reads compatible with the first groups
HOMZ: 1 for homozygous ACGT calls, 0 for heterozygous or deletion
DEL1: 1 for heterozygous deletions, 0 for all others
DEL2: 1 for homozygous deletions, 0 for all others
EDIST0: 1 if the nucletotide is the first or last of the region, 0 otherwise
EDIST1: 1 if the nucletotide is 1 position from the start or the end of the
region, 0 otherwise
EDIST2: 1 if the nucletotide is 2 positions from the start or the end of the
region, 0 otherwise
```

```
EDIST: the distance of the nucleotoide from the region start or end (whichever is smaller)
HAS_ALT: 1 if more than one group, 0 otherwise
MANY_ALT: 1 if more than two groups, 0 otherwise
ALT_USED: 1 if 2 groups are used for call, 0 otherwise
DIST1: (COV – GLOBAL_COV) / sqrt(GLOBAL_COV)
DIST2: ((max(A,C,G,T,GAP) – 0.5 * (A + C + G + T + GAP)) / sqrt(COV)) * ALT_USED
IGNORE: 1 if some groups were ignored  for calling, 0 otherwise
RATIO: max(A,C,G,T,GAP) / (A + C + G + T + GAP) * (GRP != 1)
SUB: Nucleotide position in insertion
```

Formulas with boolean operators use standard C semantics (true = 1, false = 0).

**Model 1 – call selection**
**Diploid**

| | Estimate |
|---|---|
| Intercept | 1.549817e+01 |
| COMP_2/G0_COMP | 3.214268e+00 |
| HOMZ | -1.603723e+01 |
| DEL1 | 4.057173e+00 |
| DEL2 | -1.295838e+01 |
| GLOBAL_COV | 3.327203e-01 |
| EDIST0 | -2.055305e+00 |
| EDIST1 | -1.914959e+00 |
| EDIST2 | -5.105844e-01 |
| EDIST | 5.987854e-02 |
| HAS_ALT | -7.634908e-01 |
| DIST1 | 1.563516e+00 |
| DIST2 | -1.233070e+01 |
| (DIST1 + 0.5) * (DIST1 > -0.5) | -3.456876e-01 |
| (DIST1 - 2) * (DIST1 > 2) | -1.089758e-01 |
| (DIST1 - 3) * (DIST1 > 3) | -8.686674e-01 |
| DIST2$^2$ | -6.547970e-01 |
| G0_COMP/GLOBAL_COV | -1.655326e+00 |
| (G0_COMP/GLOBAL_COV)$^2$ | 2.113226e-01 |
| (EDIST - 40)$^3$ * (EDIST > 40) | 2.992796e-03 |
| (EDIST - 45)$^3$ * (EDIST > 45) | -6.197973e-03 |
| IGNORE | -2.224370e-01 |
| RATIO | -1.255600e+02 |
| RATIO$^2$ | 3.233437e+02 |
| RATIO$^3$ | -2.755079e+02 |
| RATIO$^4$ | 7.897496e+01 |
| EDIST$^2$ | -8.887499e-04 |
| HOMZ * (EDIST <= 5) | -2.998684e-01 |
| COMP_2 / G0_COMP) * GLOBAL_COV | -1.062955e-01 |
| HOMZ * GLOBAL_COV | -2.855130e-01 |
| DEL1 * GLOBAL_COV | -9.098014e-02 |
| DEL2 * GLOBAL_COV | -2.018754e-01 |
| DEL2 * EDIST | 7.388170e-02 |
| DEL2 * HAS_ALT | -4.950726e+00 |
| DEL2 * DIST1 | -6.573440e-01 |
| DEL2 * DIST2 | 1.337017e+01 |
| HOMZ * DIST2$^2$ | 2.234410e+00 |
| HOMZ * G0_COMP / GLOBAL_COV | 2.994476e+00 |
| HOMZ * (G0_COMP/GLOBAL_COV)$^2$ | -4.286640e-01 |
| HOMZ * DIST1 | -8.026551e-01 |
| HOMZ * DIST2 | 9.614824e+00 |
| DEL1 * EDIST | -1.301157e-01 |
| EDIST * DIST1 | -1.017782e-02 |
| DIST1 * EDIST$^2$ | 1.413317e-04 |
| DEL1 * EDIST$^2$ | 2.472375e-03 |

### Haploid

| | Estimate |
|---|---|
| Intercept | 2.734031375 |
| EDIST0 | -8.395304525 |
| (EDIST == 1) OR (EDIST == 2) | -2.292773866 |
| (EDIST - 45) * (EDIST > 45) | 1.502826728 |
| DIST1 | 0.617528244 |
| EDIST * DIST1 | -0.009752782 |

### Model1B – quality score

### Diploid

| | |
|---|---|
| Intercept | 5.625990e+00 |
| HOMZ | -1.926639e+00 |
| DEL2 | -4.149465e+00 |
| DIST1 | 1.976799e+00 |
| DEL1 | -3.674773e-01 |
| GLOBAL_COV | 2.505259e-01 |
| COMP_2 / G0_COMP | 3.530792e+00 |
| MANY_ALT | 2.384205e-01 |
| HAS_ALT | -1.893987e+00 |
| EDIST0 | 2.488365e+00 |
| EDIST1 | 3.614451e+00 |
| EDIST2 | -8.343540e-01 |
| (DIST1 + 2) * (DIST1 > (-2)) | -3.608020e-01 |
| (DIST1 - 2) * (DIST1 > (2)) | -1.369033e+00 |
| DIST2 | -8.717219e-01 |
| (DIST1 + 1) * (DIST1 > (-1)) | -5.990449e-01 |
| G0_COMP / GLOBAL_COV | -5.090870e-01 |
| (EDIST - 35) * (EDIST > 35) | 7.200000e-02 |
| (EDIST - 30) * (EDIST > 30) | -6.277709e-02 |
| (EDIST - 45) * (EDIST > 45) | 1.407460e-01 |
| $GLOBAL\_COV^2$ | -3.807892e-03 |
| IGNORE | -5.524936e-01 |
| SUB > 0 | -1.085515e+00 |
| HOMZ * (EDIST < 5) | 1.155368e+00 |
| RATIO | -1.489082e+02 |
| $RATIO^2$ | 6.542650e+02 |
| $RATIO^3$ | -9.392902e+02 |
| $RATIO^4$ | 4.360459e+02 |
| DIST1 * DEL1 | -2.069432e-01 |
| HOMZ * GLOBAL_COV | 1.598539e-01 |
| DEL2 * GLOBAL_COV | 2.304383e-01 |
| DIST1 * GLOBAL_COV | -1.981619e-02 |
| DEL1 * GLOBAL_COV | 5.554233e-02 |
| DEL2 * HAS_ALT | -5.609686e-01 |
| DEL2 * DIST1 | 7.001617e-01 |
| DEL1 * $DIST1^2$ | 1.859963e-01 |
| HOMZ * DIST1 | 6.971654e-01 |
| HOMZ * DIST2 | -1.003972e-01 |
| HOMZ * G0_COMP / GLOBAL_COV | -6.196470e-01 |
| HOMZ * $(G0\_COMP / GLOBAL\_COV)^2$ | 1.267673e-01 |
| GLOBAL_COV * (SUB > 0) | -1.475575e-01 |
| DIST1 * (SUB > 0) | -7.022790e-01 |
| HOMZ * $DIST2^2$ | 2.281341e-01 |
| DIST1 * DEL1 * GLOBAL_COV | 1.536606e-02 |
| HOMZ * DIST2 * (SUB > 0) | -5.997786e-01 |

### Haploid

| | Estimate |
|---|---|
| Intercept | 7.7911387 |

| | |
|---|---|
| (EDIST - 45) * (EDIST > 45) | 0.7390936 |
| SUB > 0 | -5.7026205 |
| HOMZ * (EDIST < 5) | -0.9447409 |

**Model 2 – the quality of reference calls**

| | Estimate |
|---|---|
| Intercept | -7.339851e+00 |
| DIST1 | 2.457963e+00 |
| $DIST1^2$ | -2.092731e-01 |
| $DIST1^3$ | 1.757365e-02 |
| EDIST | 1.174253e+00 |
| COV | 2.189787e-01 |
| GLOBAL_COV | 7.489705e-01 |
| $COV^2$ | -1.873808e-02 |
| $COV^3$ | 2.716039e-04 |
| (DIST1 + 0.5) * (DIST1 > -0.5) | -5.814003e-01 |
| (DIST1 - 3) * (DIST1 > 3) | -8.967198e-02 |
| EDIST2 | 1.881940e+00 |
| $EDIST^2$ | -1.146688e-01 |
| $EDIST^3$ | 4.807719e-03 |
| $EDIST^4$ | -9.036972e-05 |
| $EDIST^5$ | 6.263128e-07 |
| DIST1 * EDIST | -4.384856e-03 |
| COV * GLOBAL_COV | -3.309976e-02 |
| GLOBAL_COV * $COV^2$ | 9.086561e-04 |
| GLOBAL_COV * $COV^3$ | -9.727565e-06 |
| EDIST * GLOBAL_COV | -9.141201e-05 |
