## Supplementary file 4: Diploid model for "KATK: fast genotyping of rare variants directly from unmapped sequencing reads"

### Probability to observe a given vector of frequencies

The number of reads containing in a given genomic position letter A, T, C, G and gap are denoted respectively by $n_A, n_T, n_C, n_G, n_{gap}$. The total number of reads covering one position is $n = n_A + n_T + n_C + n_G + n_{gap}$.

The probability to observe a set of frequencies given the genotype ($geno$) and the total number of reads covering one genomic position ($n$) can be calculated by using multinomial distribution:

$$P(n_A, n_T, n_C, n_G, n_{gap}|geno) = \frac{n!\, p_{A|geno}^{n_A}\, p_{T|geno}^{n_T}\, p_{C|geno}^{n_C}\, p_{G|geno}^{n_G}\, p_{gap|geno}^{n_{gap}}}{!n_A!n_T!n_C!n_G!n_{gap}},$$

where $p_{A|geno}$ is the conditional probability to observe a read with letter A in particular genomic position given the true genotype is $geno$, $p_{C|geno}$ is the conditional probability to observe a read with letter C in the same position given the true genotype is $geno$ etc. If the true genotype is homozygous (for example AA), then the conditional probabilities to observe any other character (C, T, G, gap) is equal to $\varepsilon/4$, where $\varepsilon$ denotes the probability of sequencing error. For example:

$$\begin{array}{rlrll}
p_{C|geno=AA} &= \varepsilon/4 & p_{A|geno=CC} &= \varepsilon/4 & \ldots \\
p_{G|geno=AA} &= \varepsilon/4 & p_{G|geno=CC} &= \varepsilon/4 & \\
p_{T|geno=AA} &= \varepsilon/4 & & \ldots & \\
p_{gap|geno=AA} &= \varepsilon/4 & & &
\end{array}$$

For homozygous genotypes the probability of correct reading is $1-\varepsilon$, so $P_{A|geno=AA} = 1-\varepsilon$, $P_{C|geno=CC} = 1-\varepsilon$ etc.

For heterozygous genotypes the probabilities to observe a read with wrong letter in the position of interest are still equal to $\varepsilon/4$: $p_{T|geno=AC} = \varepsilon/4$, $p_{G|geno=AC} = \varepsilon/4$ etc.

But as there are two possibilities for correct reads the conditional probabilities for letters present in the true genotype are equal to $(1-\varepsilon)/2+\varepsilon/8$. Comment: If the true genotype is AC, then one may see the letter A either because the read should contain A and it is correctly sequenced as A or because the read should contain C but C is misread as A. Due to the latter possibility (one makes sequencing error but the misread letter is actually present in the genotype) one has to add the $\varepsilon/8$ to the conditional probability.

Therefore the conditional probabilities to observe one particular letter present in the true genotype in a given read for heterozygous genotypes are taken to be $p_{A|geno=AC} = (1-\varepsilon)/2 + \varepsilon/8 = 1/2 - 3/8\varepsilon$, $p_{C|geno=AC} = 1/2 - 3/8\varepsilon$, $p_{A|geno=AG} = 1/2 - 3/8\varepsilon$, etc.
