## Supplementary text 2: Simulation parameters for "KATK: fast genotyping of rare variants directly from unmapped sequencing reads"

**Simulated genomes**

To test the performance, sensitivity and selectivity of KATK at different sequencing coverages we created 20 sets of simulated reads, 10 male and 10 female. The coverages varied from 5 to 50 with a step of 5. Read were generated from the human reference genome (GRCh38p10) with wgsim (https://github.com/lh3/wgsim) separately for autosomes, sex chromosomes and mitochondrion and then mixed together. Read length was 151, error rate 0.005, mutation rate 0.0015 and the number of mitochondrion genomes per cell 100. The generated FASTQ files of relevant chromosomes were combined and the list of mutations was used for testing KATK sensitivity.
